# Complementary genetic and epigenetic changes facilitate rapid adaptation to multiple global change stressors

**DOI:** 10.1101/2024.03.20.585843

**Authors:** Reid S. Brennan, James A. deMayo, Michael Finiguerra, Hannes Baumann, Hans G. Dam, Melissa H. Pespeni

## Abstract

To persist in the geologically unprecedented rates of global change, populations can adapt or acclimate. However, how these mechanisms of resilience interact, particularly the role of epigenetic variation in long-term adaptation, is unknown. To address this gap, we experimentally evolved the foundational marine copepod *Acartia tonsa* for 25 generations under ocean acidification, warming, their combination, and control conditions then measured epigenomic, genomic, and transcriptomic responses. We observed clear and consistent epigenomic and genomic divergence between treatments, with epigenomic divergence concentrated in genes related to stress response and the regulation of transposable elements. However, epigenetic and genetic changes occurred in different regions of the genome such that regions with significant methylation divergence had 2-2.5 fold lower F_ST_ than regions without methylation divergence. This negative relationship between epigenetic and genetic divergence could be driven by local inhibition of one another or distinct functional targets of selection. In contrast, epigenetic divergence was positively linked to gene expression divergence, indicating that epigenetic changes may facilitate phenotypic change. Taken together, these results suggest that unique, complementary genetic and epigenetic mechanisms promote resilience to global change.

**Significance Statement:** Organisms must adapt or acclimate to survive global change, but how these processes interact and the role of epigenetic variation is unknown. To address these gaps, we experimentally evolved the marine copepod *Acartia tonsa* for 25 generations in global change conditions and measured their genomic, epigenomic, and gene expression responses. We found that both genetic and epigenetic changes contributed to resilience and were inversely related, acting in different regions of the genome. Epigenetic changes were functionally linked to the regulation of stress and transposable elements and correlated with shifts in gene expression. Therefore, the resilience of populations to ongoing global change is driven by the complementary contribution of both genetic and epigenetic mechanisms.

## Introduction

Global conditions are changing at a geologically unprecedented rate (1). To persist in a given location, a population can respond using two mechanisms: plasticity and adaptation. Plasticity allows an organism to adjust its physiology in response to environmental stress (2), with epigenetic modifications playing a crucial role in facilitating these adjustments (3). In fact, recent studies across a broad range of taxa have shown that these epigenetic changes, along with the resulting phenotypic traits, can be inherited across generations (4). A more well understood mechanism for rapid adaptation across generations is selection on standing genetic variation (5). However, no studies to date have simultaneously measured genetic, epigenetic, and transcriptomic responses during adaptation to global change (6) despite the urgent need and call for such studies (7, 8). Here, we measure epigenomic, transcriptomic, and genomic responses of copepod populations experimentally evolved in ocean warming, ocean acidification, and combined ocean warming and acidification conditions to assess the interplay between these unique response mechanisms during rapid global change adaptation.

Plasticity and adaptation may interact in complex ways. Phenotypic plasticity allows individual genotypes to shift their phenotypes to match the environment and maximize fitness. However, for most populations, plasticity alone will be insufficient to completely buffer the effects of global change (9) and rapid adaptation or evolutionary rescue (10) from standing genetic variation will be crucial (11). These two mechanisms could facilitate or inhibit one another during responses to changing environments (12). For example, plasticity can facilitate short-term population persistence thereby allowing time for selection to act and adaptation to proceed (13). Alternatively, when plasticity moves a phenotype closer to the environmental optimum, it may effectively shield a population from selection thereby inhibiting adaptation (2).

Epigenetic mechanisms are key to understanding the interaction between plasticity and adaptation during evolutionary rescue. Epigenetic variation is broadly defined as changes that affect gene expression, such as DNA methylation or chromatin modification, that do not involve changes in underlying DNA sequence (14). Because epigenetic changes impact gene expression regulation, modifications at the epigenetic level can have phenotypic effects (15). For instance, in corals, methylation correlates with gene expression, where genes with stable expression tend to have high methylation and environmentally responsive genes have low methylation (16, 17). Further, studies in model organisms have shown the inheritance of gene expression phenotypes across generations in response to acute stressors experienced in the parental generation, e.g., temperature and nutrition in *Drosophila* (18, 19), pollutants in zebrafish (20), and temperature in *C. elegans* (21). These results suggest that epigenetic patterns are generally responsive to environmental conditions and can affect (17) the ability of populations to respond to changing environmental conditions (22).

A powerful approach to understand the mechanisms that shape a species capacity for resilience in future global environments is to simulate future conditions and observe population responses across generations in real time, known as experimental evolution (23). When combined with genomic methods, it is possible to identify the genomic targets of selection as well as the epigenomic and phenotypic responses enabling evolutionary rescue (6). Here, we leveraged a novel approach integrating genomic, epigenomic, and transcriptomic data with 25 generations of experimental evolution under ambient (AM, 18°C; 400 μatm pCO_2_) and simulated future ocean acidification (OA, 18°C; 2,000 μatm pCO_2_), ocean warming (OW, 22°C; 400 μatm pCO_2_), and their combination (OWA; (22°C; 2,000 μatm pCO_2_) in the marine copepod, *Acartia tonsa* (Fig. 1). We then quantified the relationships between genomic, epigenomic, and transcriptomic changes across the treatments, providing an unprecedented view into the relationship of these mechanisms during evolutionary rescue across multiple generations. Copepods provide an ideal model as their small size (∼1 mm), short generation times (∼3 weeks), and genomic resources make multi generation and complex experiments with high replication and census sizes tractable (24). Additionally, copepods are foundational to marine ecosystems worldwide as they are one of the most abundant metazoans (25) and provide critical links between higher level consumers and primary producers (26). *Acartia tonsa* specifically is a cosmopolitan estuarine and nearshore species that is dominant in the Atlantic ocean (27) and serves as a main prey item for fishes (28). Therefore, the resilience of copepods and *A. tonsa* under global change conditions has important implications for the stability of marine ecosystems across the globe.

**Figure 1:**
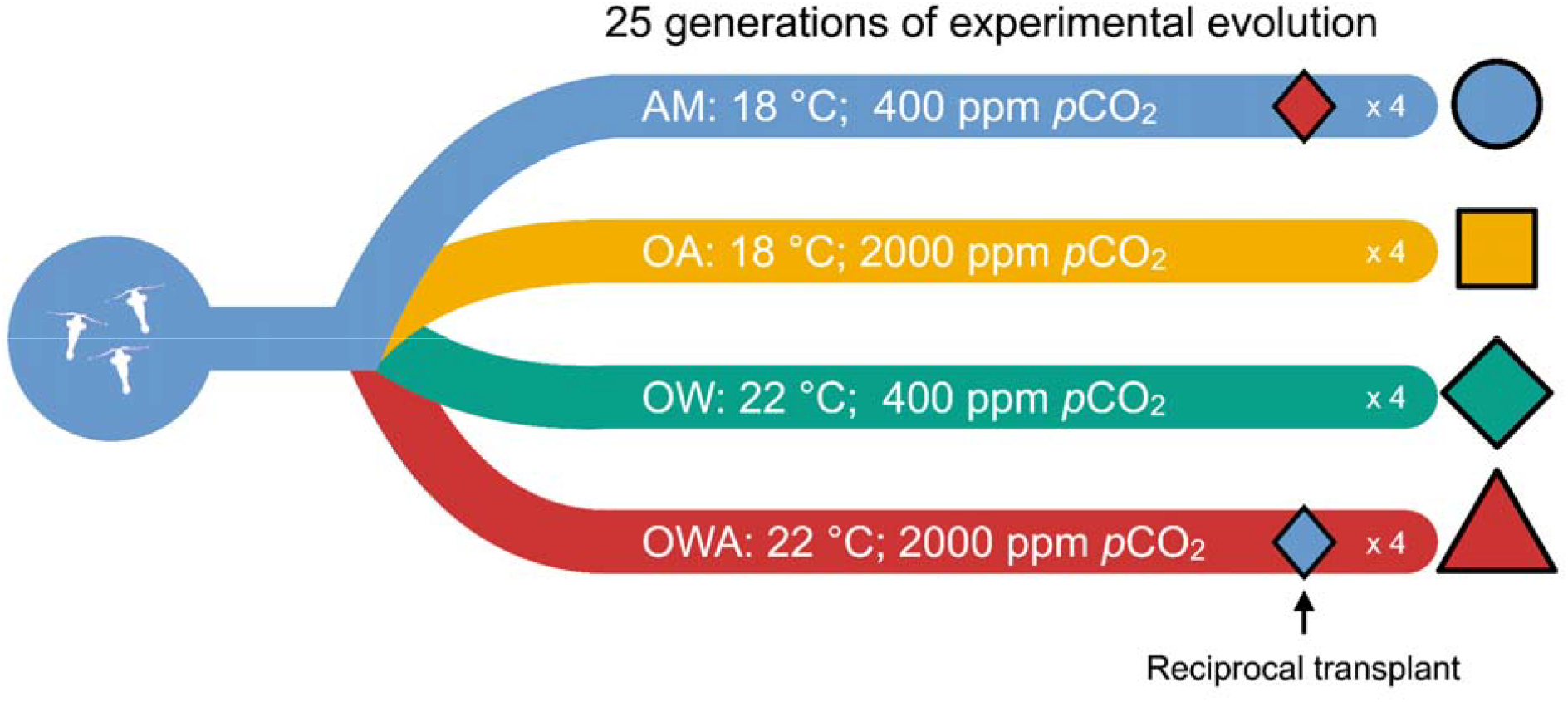
Schematic of the experimental design. The starting lab population was founded with 160 female and 80 male wild-caught adults and each treatment consisted of four replicates and census sizes of at least 3,000. The experiment was run for 25 generations, sequencing animals for genomics and epigenomics at the end. To understand potential changes in plasticity, we conducted a reciprocal transplant of AM into OWA and OWA into AM at generation F21 and quantified the transcriptomic responses.

## Results

### Genome-wide epigenetic variation

After 25 generations of evolution in ambient (AM), ocean acidification (OA), ocean warming (OW), and combined warming and acidification (OWA), methylation frequencies of pools of 30 individuals from each of four replicates per treatment were quantified using reduced representation bisulfite sequencing (RRBS). This resulted in 96,207 methylated sites sequenced across all samples at a depth of at least 15x per sample. Mean methylation percentage was different across treatments (ANOVA p-value = 0.002) and was driven by a decrease in methylation from F0 to F25 (mean methylation ± SD: F0: 0.254 ± 0.003; F25: 0.229 ± 0.009; Tukey test: p < 0.03; Fig. S1; table S1) but did not differ between treatments at F25 (p > 0.7).

Genome-wide methylation frequencies were summarized for all samples using multidimensional scaling (MDS) and generally showed clustering by treatment groups (Fig. 2A, Fig. S2). The OWA and warming treatments clustered together and were largely distinct from ambient and acidification along dimension 1 (Fig. 2A). Differential methylation analysis showed that OWA had the largest number of loci changing methylation frequency relative to ambient controls (753 loci; methylation change > 10%; FDR < 0.05) followed by warming (161 loci) and acidification (128 loci; Fig. 3). There was limited shared change in methylation between treatments with 48 loci overlapping between OWA and warming, 50 between OWA and acidification, 16 between warming and acidification, and 8 shared across all three treatments (Fig. 2B) suggesting the combined treatment results in unique, synergistic methylation changes.

**Figure 2:**
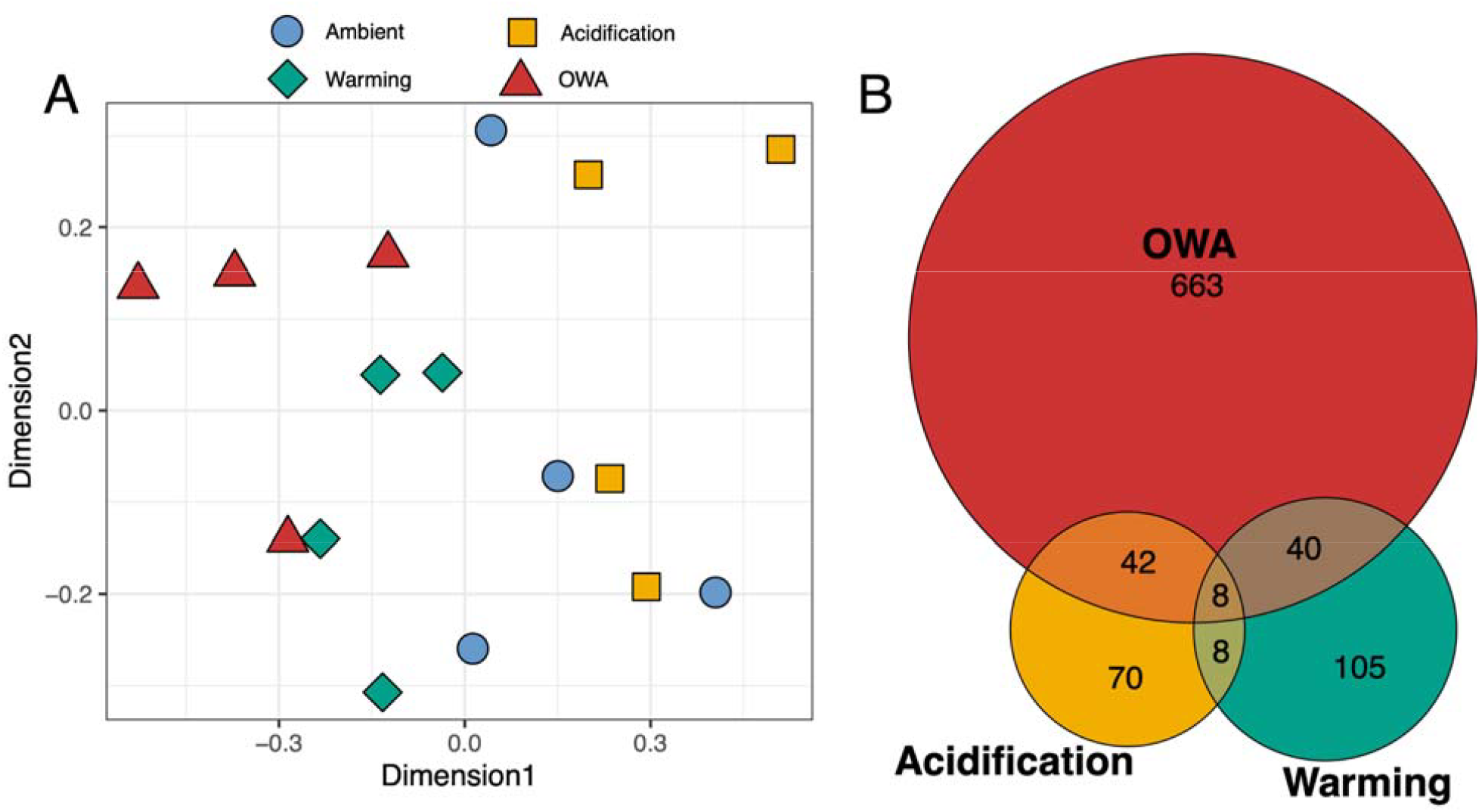
Overview of epigenetic changes. (**A)** Multidimensional scaling of methylation frequencies of 96,207 loci across replicate selection populations. **(B)** Venn diagram of significant methylation changes relative to ambient F0 and F25 (10% change in frequency, FDR < 0.05).

**Figure 3:**
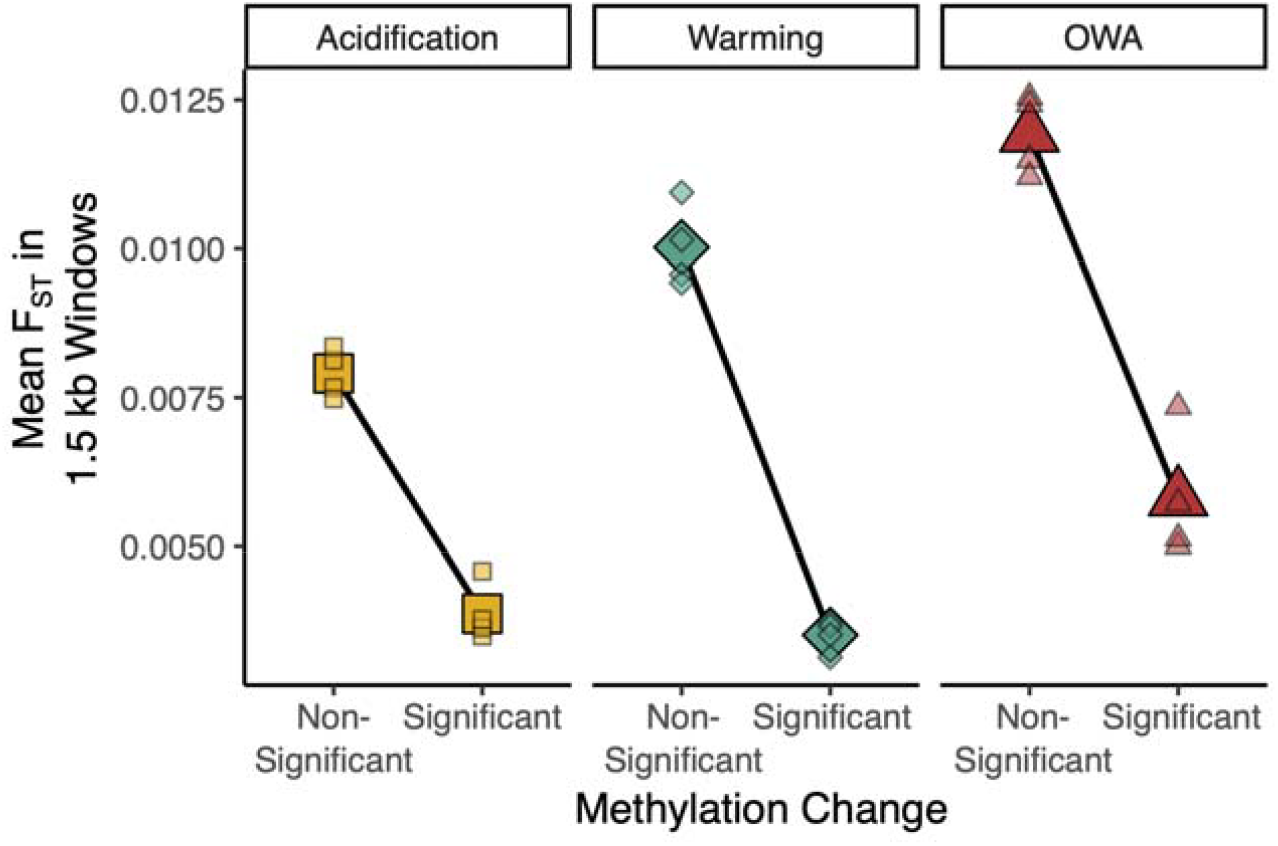
Relationship between F_ST_ and methylation change. The genome was broken into 1.5 kb windows that contained at least 5 SNPs and 5 methylation sites resulting in 910 windows across the genome. Each small point represents a replicate and the large symbols are the mean of all replicates. If a window contained at least one significant change in methylation from the ambient treatment it was considered significant. The difference in F_ST_ between windows containing significant and non-significant methylation changes is greater than would be expected by change for all treatments (P = 0.0001).

### Functional targets of epigenomic changes

The significant epigenetic changes were enriched for numerous functional categories. The largest enrichment was observed for OWA (16 terms) followed by warming (4 terms) and acidification (2 terms). Across all treatments, epigenetic changes were enriched for functions related to proteolysis (Table S2, p < 0.05, GO:0006508 and GO:0008233). Warming and OWA shared enrichment for DNA recombination (GO:0006310) while acidification and OWA were enriched for categories related to the regulation of transposable elements (GO:0006313; GO:0032197). Further, OWA was enriched for several categories related to viral components (GO:0075732, GO:0044826, GO:0075713, GO:0046718, GO:0044423).

### Interaction between epigenomic and genomic changes

Epigenetic and genetic changes were inversely related following 25 generations of selection. Allele frequencies for 394,667 loci across all samples were reported in Brennan et al. (29). Comparing allele frequency change from F0 to F25 to methylation changes summarized at the gene level showed a significant negative relationship where higher allele frequency changes corresponded to lower methylation changes (Fig. S3; p < 0.001). To directly quantify the relationship between the epigenetic and genomic changes, we compared changes in 1.5 kb windows across the genome for windows that had at least 5 SNPs and 5 epi-loci, resulting in 910 windows. There was again a negative relationship between methylation and allelic changes where windows containing significant methylation changes had lower allele frequency divergence (Fig. 3). For OWA, windows without a significant change in methylation had 2 fold higher F_ST_ (0.012) than those with (0.006; randomization test: p = 0.0001; Fig. S4, S5); similar patterns were observed for warming (Non-significant methylation change window F_ST_ = 0.010, Significant methylation change window F_ST_ = 0.004; p = 0.0001) and acidification (Non-significant methylation change window F_ST_ = 0.008, Significant methylation change window F_ST_ = 0.004; p = 0.0001). This relationship was robust to changes in window size (Fig. S6) and was present for warming and OWA, but not acidification, when calculating the reverse: the mean methylation changes for windows with significant allele frequency divergence (Fig. S7). Thus, epigenetic and genetic changes were inversely related after 25 generations.

To determine if there was a relationship between genetic variation and methylation change we quantified Tajima’s π between genomic regions with and without significant methylation changes. For the OWA line, genetic variation was higher in regions with significant methylation change than those with non-significant changes (Tajima’s π: 0.0196 ± 0.0007 vs. 0.0171 ± 0.0001; p = 0.0001; Kolmogorov-Smirnov Test; Fig. S8). There was no difference between these regions for acidification (0.0165 ± 0.0001 vs. 0.0169 ± 0.003; p = 0.2) or warming (0.0169 ± 0.0001 vs. 0.0205 ± 0.002; p = 0.7). However, the total number of windows overlapping with significant methylation changes was much lower for these two treatments than for OWA, which may make estimates unreliable (overlapping window number: acidification = 30; warming = 90; OWA = 456). These results suggest that epigenetic change is more likely in regions of the genome with higher genetic variation.

### Correlations between epigenetic changes and gene expression

There were significant relationships between epigenetic changes and gene expression, indicating potential phenotypic effects of the epigenetic changes. We previously quantified gene expression changes in the ambient and OWA lines in their treatment conditions and following a reciprocal transplant (30); we integrate these data with the current methylation data. At the gene level, divergence in expression between OWA and ambient lines was significantly associated with changes in methylation. For genes that had at least five methylation sites and one of these sites with significant methylation change, larger changes in methylation for OWA were correlated with larger divergence in gene expression (Fig. 4A; R-squared = 0.025, p = 0.015) suggesting a role for epigenetic variation in long-term divergence in gene expression in static conditions. To test for a relationship between percent methylation and plasticity in gene expression, we used the reciprocal transplant to characterize genes as plastic (significant change in expression) or not plastic (no change in expression). Genes that showed significant expression change following transplant of the OWA line to ambient conditions had significantly lower methylation than those with no change in gene expression (Fig. 4B; p= 0.02; mean methylation ± se: Non-plastic: 0.115 ± 0.005; Plastic: 0.026 ± 0.004). This result was also present for the ambient line when transplanted to OWA conditions (Fig. S9; p= 8.6e-8; mean methylation ± se: Non-plastic: 0.133 ± 0.006; Plastic: 0.068 ± 0.0070). Taken together, these results suggest that fixed differences in gene expression as populations adapted to global change were reinforced with changes in methylation, while genes that were plastically responsive to sudden changes in environment remained lowly methylated.

**Figure 4:**
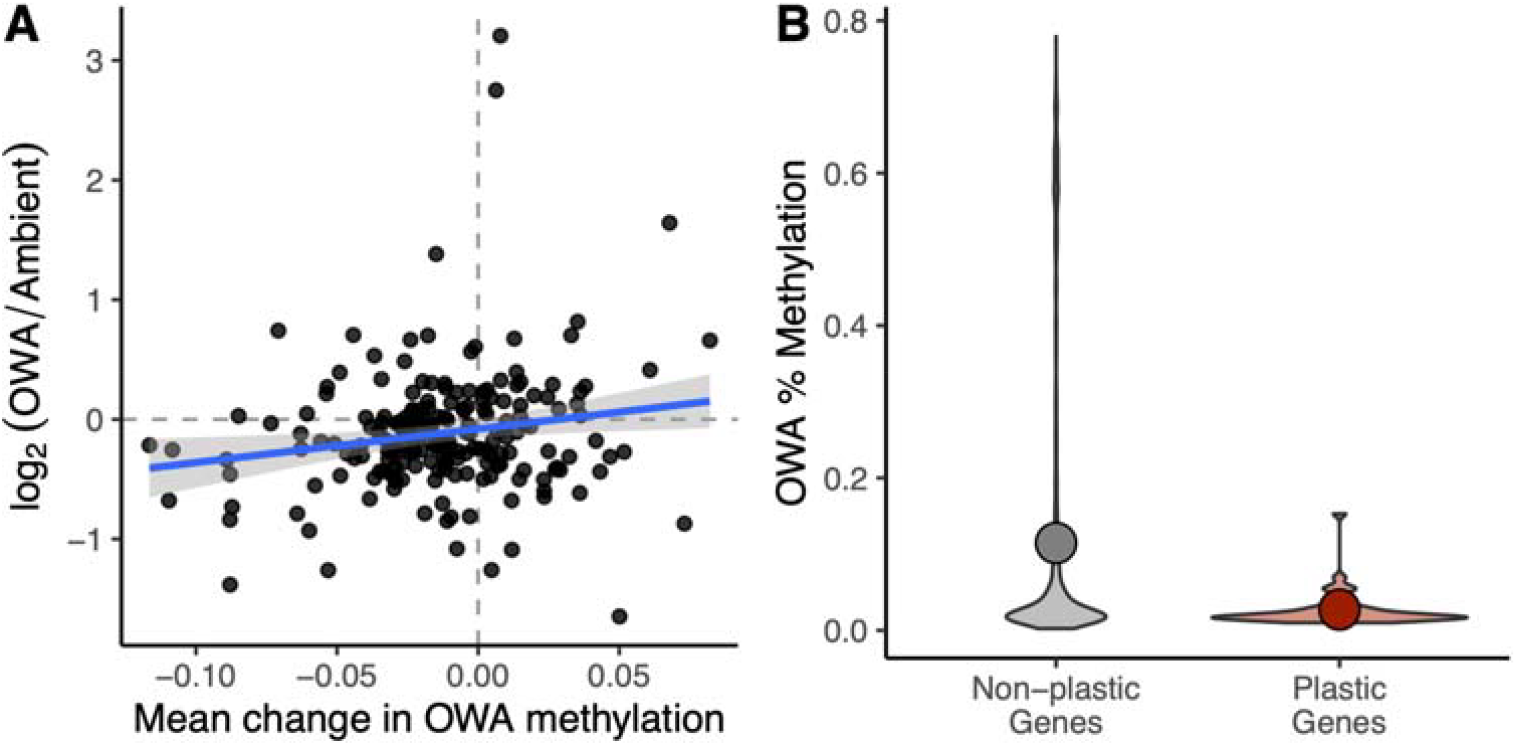
Methylation and gene expression changes. **(A)** Divergence in gene expression between OWA and ambient lines versus the mean change in OWA methylation from the ambient line. Genes were required to have at least 5 methylation sites with at least one significantly diverged methylation site in OWA. Solid blue line is the regression between these factors (R-squared = 0.025, p = 0.015). **(B)** The percent methylation of plastic vs. non-plastic genes following transplant of the OWA line from OWA to ambient at generation F21. Solid points show the mean methylation percent for each group (Non-plastic: 0.115 ± 0.005; Plastic: 0.026 ± 0.004; mean ± se). Non-plastic genes had significantly higher methylation percent than plastic genes (p= 0.019, Kolmogorov-Smirnov Test)

## Discussion

The interaction between plasticity and adaptation is an integral component underlying the resilience, or lack thereof, of populations to rapidly changing environments. While how these components interact during adaptation across natural environmental gradients has been studied (17, 31, 32), limited work has focused on how epigenetic and genetic changes might drive phenotypic responses during evolutionary rescue from global change (7, 33). Our results show that epigenetic and genetic changes are negatively related and affect unique regions of the genome (Fig. 3). Further, epigenetic divergence was positively correlated with gene expression divergence in the lineages adapted to global change stressors (Fig. 4). Thus, genetic and epigenetic changes act in a complementary manner to facilitate physiological resilience during evolutionary rescue from global change.

The negative relationship between epigenetic and genetic changes indicates that these mechanisms function in unique regions of the genome or may inhibit one another locally (Fig. 3). Similar results have been found in three-spined stickleback, *Gasterosteus aculeatus*, where adaptive changes in methylation were more likely to occur in regions of low genetic divergence, which the authors suggest is due to the epigenetic modifications achieving the phenotypic effect in place of genetic changes (34). That is, if functional changes are achieved via epigenetic shifts, this might preclude selection from acting on genetic variation and thus reduce genetic divergence in these regions. In addition, we found that regions with methylation divergence also harbored higher levels of nucleotide diversity relative to regions without methylation divergence (Fig. S5). Similar patterns of increased genetic diversity associated with methylation divergence have been found in *G. aculeatus* that have adapted from marine to freshwater environments (31), though these changes occurred over much longer timescales, around 700 years, ∼230-700 generations (35), relative to the 25 generations in the current study. The authors suggest that environmentally responsive methylation sites might be under relaxed selection at the genetic level, thus increasing genetic diversity in these regions. Alternatively, higher levels of methylation can influence mutation rates with a higher probability of spontaneous deamination of methylated cytosines to thymines (36, 37). In our 25-generation study, the combination of reduced genetic divergence in the epigenetically responsive regions of the genome and the positive relationship between epigenetic and transcriptomic divergence support the hypothesis that epigenetic changes can functionally replace genetic changes locally within the genome during short-term responses to the environment (38), leading to increased genetic variation and decreased genetic responses to selection in these regions.

The observed epigenetic changes likely have adaptive phenotypic consequences. At the gene level, we found significant functional enrichment across all treatments related to proteolysis. Proteolysis is an adaptive response to stress as it removes damaged proteins (39). Other processes were related to the regulation of transposable elements (TEs), particularly under OWA conditions. The activation of TEs under stress has been understood since McClintock’s classic work (40, 41) and has since been found in numerous systems (40, 42) to influence gene expression both in beneficial and detrimental ways (43). Under rapid environmental change, including global change, the regulation of transposable elements can be adaptive and help organisms match their phenotype to the environment (44) and this process can be dictated by epigenetic mechanisms (42, 45). Together with our results, these studies suggest that global change stress can be countered by the epigenetic regulation and expression of transposable elements, which can contribute to obtaining an adaptive phenotype under these conditions.

In addition to functional enrichment, there was a positive correlation between changes in methylation and gene expression, suggesting that epigenetic divergence was linked to phenotypic divergence between the selection regimes (Fig. 4). Indeed, methylation levels have been consistently linked to gene expression in diverse organisms, including during responses to global change conditions (17, 46–51) and in other copepods (52). Together, these results suggest that epigenetic responses may help to facilitate physiological resilience to global change conditions. We previously found a rapid recovery of fitness in the OWA line after only three generations (53) coupled with a loss of plasticity and fitness with transplantation at the 21st generation (30). While there was a strong signal of adaptation in later generations (29), the genetic and epigenetic changes at early generations were not quantified. We hypothesize that plastic epigenetic shifts may in part underlie the immediate fitness recovery and allow population persistence and time for adaptation to proceed, reminiscent of the Baldwin Effect (13); additional data focusing on short-term responses to OWA conditions would be required to test this hypothesis.

The fate of the epigenetic changes observed here over longer timescales remains unclear. In studies focused on longer-term divergence and adaptation between environments, it is common that genetic and epigenetic divergence are positively associated (17, 54). For example, in the coral, *Acropora millepora*, methylation patterns that are fixed between populations (i.e., not responsive to the environment) occur in regions with elevated genetic divergence (17). This suggests that under stable conditions, plastic epigenetic changes may become fixed due to underlying genetic changes, i.e., genetic assimilation (55, 56). Similarly, a study in *Arabidopsis thaliana* showed a positive relationship between epigenetic and gene expression divergence among latitudinally adapted accessions (57). Over the 25 generations in the current experiment, it is likely that there has been insufficient time for genetic divergence to fix the previously plastic epigenetic response. Future studies should combine short and long-term experiments to determine the pace at which epigenetic variation becomes assimilated due to genetic divergence.

While there were observed links between methylation, genetic, and transcriptomic changes, there were limitations to the study that should be taken into account. First, the method used to quantify epigenetic variation was a reduced representation approach (RRBS), which covers only a portion of the genome. Full genome quantification of methylation markers would provide a higher resolution view of the relationship between genetic and epigenetic changes. Second, we do not know if the epigenetic changes between lines are responsive to the environment or fixed between populations. To understand this aspect of the data, it would be necessary to conduct transplant experiments and quantify methylation levels following transplant across generations. Similarly, it is unknown the order in which genetic and epigenetic changes arose in the lines, which would require fine-scale quantification of both factors across generations at the early stage of the experimental evolution, which was not done here. Though we found consistent, directional epigenetic change after 25 generations of selection, it is unknown if these changes are due to transgenerational inheritance of epigenetic states or re-establishment of epigenetic marks at each generation in response to the developmental environment. Importantly, due to the negative relationship between genetic and epigenetic changes, epigenetic differences among groups are unlikely to be due to linked selection on genetic loci. Understanding the fine-scale interactions between epigenetic and genetic changes during the early stages of adaptation to global change conditions will require additional experiments and is critical to understand how these factors promote resilience to ongoing environmental change.

## Materials and Methods

### Data generation

The details of the experiment have been reported previously in Dam et al., (53). Adult copepods (n=1000) were collected from Esker Point Beach in Groton, Connecticut, USA (41.320725° N, 72.001643° W) in June 2016 and common gardened for three generations. For the selection experiment, replicate cultures were started with 160 females and 80 males in each of the following conditions: ambient (18 °C, 400 μatm CO_2_, pH ∼8.2); acidification (18 °C, 2,000 μatm CO_2_, pH ∼7.5); warming (22 °C, 400 μatm CO_2_, pH∼8.2); OWA (22 °C, 2,000 μatm CO_2_, pH∼7.5). All cultures were held at a salinity of 32.75 (CI[32.4, 33.46]) Practical Salinity Units and 12 h light, 12 h dark and fed a combination of *Tetraselmis sp*., *Rhodomonas sp*., and *Thalassiosira weissflogii* every 2 to 3 d at food-replete condition (≥ 800 μg carbon L ^-1^) (58). Cultures were held in these conditions for 25, non-overlapping, generations and populations sizes were > 3,000 per replicate.

### Epigenetics library prep/sequencing

Reduced representation bisulfite sequencing was used to quantify methylation frequencies for each sample following vonHoldt et al. (59). This approach is based on a modified NEBNext Ultra library preparation that includes a restriction digest and bisulfite conversion. For each sample, 250ng of high quality genomic DNA was digested with the restriction enzyme *Msp1*, 3’ ends were repaired with adenine and Illumina methylated sequence adapters for NEBNext Ultra library preparation were ligated (New England Biolabs, Ipswich, MA, USA). To quantify bisulfite conversion efficiency, 1ng unmethylated lambda DNA (Promega) was added to each sample. Libraries were size-selected to retain fragments between 100 and 500bp. Following size selection, libraries were bisulfite converted using a Qiagen Epitech kit with the protocol for low DNA concentrations and an extended incubation period of 10 hours to ensure high conversion efficiency of unmethylated cytosines to uracils. Libraries were then PCR amplified for 15 cycles where uracil was converted to thymine through use of EpiMark Hot Start Taq (New England Biolabs, Ipswich, MA, USA) and a unique barcode was added to each sample. Libraries were pooled at equimolar concentrations and sequenced on 2 lanes of an Illumina HiSeq 4000 with a 20% phiX spike in to increase complexity and improve data quality.

### Epigenetic data analysis

Raw reads were trimmed with Trimmomatic v0.36 for quality and adapter contamination. Mapping was conducted using Bismark v0.22.3 (60) and Bowtie2 v2.2.6 with local alignments (61). Each of the two lanes were mapped separately to the *A. tonsa* genome (62) and reads with no alignment were re-mapped as single end reads to improve coverage. These data were checked for methylation bias and this bias removed when extracting CpG methylation calls with Bismark’s methylation extractor; bias was present on the first and last 2 basepairs of reads, a common pattern in RRBS data. Following methylation extraction, extracted methylation data was concatenated by sample and output to bedGraph format with the bismark2bedGraph command.

### Methylation conversion efficiency

We used the rate of methylation conversion in the spiked-in 1ng unmethylated lambda DNA to determine methylation conversion efficiency. During bisulfite conversion, unmethylated cytosine is converted to thymine while methylated cytosine remains cytosine. Because the lambda DNA is completely unmethylated, all cytosine bases should be converted to thymine during bisulfite conversion. The proportion of thymine to cytosine represents the conversion efficiency. Raw reads were aligned to the lambda genome (NCBI accession: NC_001422.1) in the same manner as to the *A. tonsa* genome and methylation calls were extracted. From this, one minus the methylation rate is the bisulfite conversion efficiency.

### Differential methylation

Bisulfite sequencing produced an average of 27.8 million paired end reads per sample. Overall mean mapping rate of trimmed sequence reads was 54 ± 14% (mean ± SD) and bisulfite conversion efficiency calculated from the lambda dna spike-in was 98.4 ± 0.3%; see table S1 for a summary of sequencing results.

Differentially methylated regions were identified in R version 3.6.0 (63) using edgeR following the methods described in Chen et al. (64). The approach treats the counts of methylated and unmethylated reads as separate observations and allows for the use of edgeR’s generalized linear model framework to incorporate complex experimental designs. We required a minimum coverage of 15x for all samples and sites with coverage greater than the 97.5% quantile (974x) were removed to control for multi-mapping regions. This filtering resulted in 96,207 methylation sites with a mean coverage of 124x and median of 77x (Fig. S10). We fit a negative binomial generalized linear model to detect differentially methylated regions between each treatment and the ambient control, considering methylation frequencies at (F0 and F25). A Benjamini-Hochberg correction was applied to control for multiple testing and differential methylation was considered significant at p < 0.05 and with methylation changes > 10 %.

### Functional enrichment

Functional enrichment was run using the weight01 method in topGO v. 2.36.0 (65, 66), requiring a minimum of five annotated genes per GO term. The background gene set included all genes assigned to methylation sites, both genic and intergenic. Of the 43,442 methylation sites assigned to genes (45% of total methylation sites), 95% were genic (41,152 loci). For enrichment tests, a gene was considered significant if it was assigned at least one loci that passed the multiple testing and 10% change in frequency thresholds.

### DNA data collection and analysis

We have previously reported results from DNA sequencing of all lines and replicates at generation F0 and F25 (29). Pools of 50 individuals were sequenced from each replicate using a sequence capture approach targeting genetic and regulatory regions (32,413 probes). After mapping to the reference genome (62), this approach resulted in 394,667 variant sites; see Brennan et al. (29) for full details. F_ST_ was estimated on a per-site level using the R package *poolfstat* (67), mean F_ST_ from the ambient line was calculated in 1.5 kb windows and the intersection between methylation sites and F_ST_ windows was identified using BEDTools (68), keeping only windows with at least 5 SNPs and 5 methylation sites. The F_ST_ distributions were then compared between windows containing a significant change in methylation, as identified in the edgeR analysis above, versus those with no change in methylation. The results were robust to both window size (1.5 vs. 5 kb) and the number of SNP or methylation sites required (3 vs. 4 vs. 5). The significance of the F_ST_ differences between groups was calculated using a randomization test with 10,000 permutations, where window significance was randomly assigned and the observed difference in mean F_ST_ was calculated. P-values were calculated as the proportion of permuted values greater than the observed value. This analysis was also run for mean methylation in 1.5 kb windows and comparing the change in methylation from F0 to F25 in windows with or without significant allele frequency divergence (from (29)).

Finally, Tajima’s π was estimated in 100bp windows using Popoolation2 (69) and the overlap between these windows and methylation sites was identified using BEDTools. Distributions of Tajima’s π were compared between windows with significant methylation change versus those without using Kolmogorov-Smirnov Tests with a Bonferroni correction in R.

### Gene expression data collection and analysis

We previously reported gene expression results from a reciprocal transplant of ambient to OWA and OWA to ambient at generation F21 (30). Gene expression was analyzed at the gene level using Salmon v0.10.2 (70) and filtered for abundance (fewer than 10 counts in >90% of the samples), resulting in 23,324 genes. Expression levels and differences were analyzed using DESeq2 (71) and plastic genes were identified as those changing expression (adjusted p-value < 0.05) within a line following transplant to the opposite environment. Differential expression between OWA and ambient lines in their home environments was also identified (adjusted p-value < 0.05).

To compare gene expression with methylation, only genes with at least five methylation sites were used. We first regressed mean methylation change with gene expression divergence to test if genes with significant methylation change in OWA relative to ambient were correlated with gene expression divergence between the lines. We next compared the mean methylation percentage for genes that were plastic versus non-plastic in their expression and compared these distributions using Kolmogorov-Smirnov Tests.

## Acknowledgements

We thank Carol Lee, Maren Wellenreuther, and Dafni Anastasiadi for helpful discussions and recommendations to improve this manuscript and Till Bayer for assistance linking genes to their GO terms for functional enrichment.

## Author contributions

Conceptualization: RSB, JAD, MF, HB, HGD, MHP

Methodology: RSB, JAD, MF, HB, HGD, MHP

Investigation: RSB, JAD

Visualization: RSB

Writing—original draft: RSB, MHP

Writing—review & editing: RSB, JAD, MF, HB, HGD, MHP

## Data and materials availability

Sequencing data are available on NCBI under BioProject accession PRJNA590963. Data files and supplemental tables can be found on Zenodo: https://doi.org/10.5281/zenodo.10797734. Code to run all analysis and produce all figures can be found on Zenodo: https://doi.org/10.5281/zenodo.10829107.

## References

1. H. O. Pörtner, et al., “Climate change 2022: impacts, adaptation and vulnerability” (IPCC, 2022).

2. S. C. Stearns, The evolutionary significance of phenotypic plasticity. Bioscience (1989).

3. D. Romero-Mujalli, et al., Emergence of phenotypic plasticity through epigenetic mechanisms. Evol. Lett. 8, 561–574 (2024).

4. D. Anastasiadi, C. J. Venney, L. Bernatchez, M. Wellenreuther, Epigenetic inheritance and reproductive mode in plants and animals. Trends Ecol. Evol. 36, 1124–1140 (2021).

5. R. D. H. Barrett, D. Schluter, Adaptation from standing genetic variation. Trends Ecol. Evol. 23, 38– 44 (2008).

6. M. A. Santos, et al., Experimental evolution in a warming world: The omics era. Mol. Biol. Evol. 41, msae148 (2024).

7. L. Bernatchez, A.-L. Ferchaud, C. S. Berger, C. J. Venney, A. Xuereb, Genomics for monitoring and understanding species responses to global climate change. Nat. Rev. Genet. 25, 165–183 (2024).

8. J. M. Eirin-Lopez, H. M. Putnam, Marine Environmental Epigenetics. Ann. Rev. Mar. Sci. 11, 335– 368 (2019).

9. A. R. Gunderson, J. H. Stillman, Plasticity in thermal tolerance has limited potential to buffer ectotherms from global warming. Proc. Biol. Sci. 282, 20150401 (2015).

10. G. Bell, Evolutionary Rescue. Annu. Rev. Ecol. Evol. Syst. 48, 605–627 (2017).

11. D. N. Reznick, J. Losos, J. Travis, From low to high gear: there has been a paradigm shift in our understanding of evolution. Ecol. Lett. 22, 233–244 (2019).

12. R. J. Fox, J. M. Donelson, C. Schunter, T. Ravasi, J.D. Gaitán-Espitia, Beyond buying time: the role of plasticity in phenotypic adaptation to rapid environmental change. Philos. Trans. R. Soc. Lond. B Biol. Sci. 374, 20180174 (2019).

13. G. G. Simpson, The Baldwin Effect. Evolution 7, 110–117 (1953).

14. D. C. H. Metzger, P. M. Schulte, Epigenomics in marine fishes. Mar. Genomics 30, 43–54 (2016).

15. K. McGuigan, A. A. Hoffmann, C.M. Sgrò, How is epigenetics predicted to contribute to climate change adaptation? What evidence do we need? Philos. Trans. R. Soc. Lond. B Biol. Sci. 376, 20200119 (2021).

16. J. L. Dimond, S. B. Roberts, Germline DNA methylation in reef corals: patterns and potential roles in response to environmental change. Mol. Ecol. 25, 1895–1904 (2016).

17. G. Dixon, Y. Liao, L. K. Bay, M. V. Matz, Role of gene body methylation in acclimatization and adaptation in a basal metazoan. Proc. Natl. Acad. Sci. U. S. A. 115, 13342–13346 (2018).

18. K.-H. Seong, D. Li, H. Shimizu, R. Nakamura, S. Ishii, Inheritance of stress-induced, ATF-2-dependent epigenetic change. Cell 145, 1049–1061 (2011).

19. A. J. Osborne, P. K. Dearden, A “phenotypic hangover”: the predictive adaptive response and multigenerational effects of altered nutrition on the transcriptome of Drosophila melanogaster. Environ. Epigenet. 3, dvx019 (2017).

20. F. Pierron, et al., Genetic and epigenetic interplay allows rapid transgenerational adaptation to metal pollution in zebrafish. Environ. Epigenet. 8, dvac022 (2022).

21. A. Klosin, E. Casas, C. Hidalgo-Carcedo, T. Vavouri, B. Lehner, Transgenerational transmission of environmental information in C. elegans. Science 356, 320–323 (2017).

22. M. Kelly, Adaptation to climate change through genetic accommodation and assimilation of plastic phenotypes. Philos. Trans. R. Soc. Lond. B Biol. Sci. 374, 20180176 (2019).

23. C. Schlötterer, R. Kofler, E. Versace, R. Tobler, S. U. Franssen, Combining experimental evolution with next-generation sequencing: a powerful tool to study adaptation from standing genetic variation. Heredity 114, 431–440 (2015).

24. H. G. Dam, Evolutionary adaptation of marine zooplankton to global change. Ann. Rev. Mar. Sci. 5, 349–370 (2013).

25. A. G. Humes, How many copepods? Hydrobiologia 292, 1–7 (1994).

26. J. Mauchline, The Biology of Calanoid Copepods (Academic Press, 1998).

27. W. Yuh Lee, B. J. McAlice, Seasonal succession and breeding cycles of three species of Acartia (Copepoda: Calanoida) in a maine estuary. Estuaries 2, 228–235 (1979).

28. J. T. Turner, “The feeding ecology of some zooplankters that are important prey items of larval fish” (NOAA Technical Report NMFS, 1984).

29. R. S. Brennan, et al., Experimental evolution reveals the synergistic genomic mechanisms of adaptation to ocean warming and acidification in a marine copepod. Proc. Natl. Acad. Sci. U. S. A. 119, e2201521119 (2022).

30. R. S. Brennan, et al., Loss of transcriptional plasticity but sustained adaptive capacity after adaptation to global change conditions in a marine copepod. Nat. Commun. 13, 1147 (2022).

31. J. Ord, T. I. Gossmann, I. Adrian-Kalchhauser, High Nucleotide Diversity Accompanies Differential DNA Methylation in Naturally Diverging Populations. Mol. Biol. Evol. 40 (2023).

32. C. F. de Carvalho, et al., DNA methylation differences between stick insect ecotypes. Mol. Ecol. (2023). 10.1111/mec.17165.

33. J. Hu, R. D. H. Barrett, Epigenetics in natural animal populations. J. Evol. Biol. 30, 1612–1632 (2017).

34. M. J. Heckwolf, et al., Two different epigenetic information channels in wild three-spined sticklebacks are involved in salinity adaptation. Sci Adv 6, eaaz1138 (2020).

35. K. Reid, M. A. Bell, K. R. Veeramah, Threespine Stickleback: A Model System For Evolutionary Genomics. Annu. Rev. Genomics Hum. Genet. 22, 357–383 (2021).

36. Y. Zhou, et al., The Impact of DNA Methylation Dynamics on the Mutation Rate During Human Germline Development. G3 10, 3337–3346 (2020).

37. J. Xia, L. Han, Z. Zhao, Investigating the relationship of DNA methylation with mutation rate and allele frequency in the human genome. BMC Genomics 13 Suppl 8, S7 (2012).

38. F. D. Klironomos, J. Berg, S. Collins, How epigenetic mutations can affect genetic evolution: model and mechanism. Bioessays 35, 571–578 (2013).

39. K. Flick, P. Kaiser, Protein degradation and the stress response. Semin. Cell Dev. Biol. 23, 515–522 (2012).

40. B. McClintock, The significance of responses of the genome to challenge. Science 226, 792–801 (1984).

41. B. Mcclintock, The origin and behavior of mutable loci in maize. Proc. Natl. Acad. Sci. U. S. A. 36, 344–355 (1950).

42. L. Schrader, J. Schmitz, The impact of transposable elements in adaptive evolution. Mol. Ecol. 28, 1537–1549 (2019).

43. V. Horváth, M. Merenciano, J. González, Revisiting the Relationship between Transposable Elements and the Eukaryotic Stress Response. Trends Genet. 33, 832–841 (2017).

44. Y.-N. Liu, R.-M. Chen, Q.-T. Pu, L. M. Nneji, Y.-B. Sun, Expression plasticity of transposable elements is highly associated with organismal re-adaptation to ancestral environments. Genome Biol. Evol. 14, evac084 (2022).

45. O. Rey, E. Danchin, M. Mirouze, C. Loot, S. Blanchet, Adaptation to global change: A transposable element-epigenetics perspective. Trends Ecol. Evol. 31, 514–526 (2016).

46. A. M. Downey-Wall, et al., Ocean Acidification Induces Subtle Shifts in Gene Expression and DNA Methylation in Mantle Tissue of the Eastern Oyster (Crassostrea virginica). Frontiers in Marine Science 7 (2020).

47. X. Dang, et al., Epigenetic-associated phenotypic plasticity of the ocean acidification-acclimated edible oyster in the mariculture environment. Mol. Ecol. 32, 412–427 (2023).

48. G. E. Hofmann, Ecological Epigenetics in Marine Metazoans. Frontiers in Marine Science 4 (2017).

49. Y. J. Liew, et al., Epigenome-associated phenotypic acclimatization to ocean acidification in a reef-building coral. Sci Adv 4, eaar8028 (2018).

50. M. R. Gavery, S. B. Roberts, Predominant intragenic methylation is associated with gene expression characteristics in a bivalve mollusc. PeerJ 1, e215 (2013).

51. J. Wan, et al., DNA methylation and gene transcription act cooperatively in driving the adaptation of a marine diatom to global change. J. Exp. Bot. 74, 4259–4276 (2023).

52. Y. H. Lee, et al., Epigenetic plasticity enables copepods to cope with ocean acidification. Nat. Clim. Chang. 12, 918–927 (2022).

53. H. G. Dam, et al., Rapid, but limited, zooplankton adaptation to simultaneous warming and acidification. Nat. Clim. Chang. 11, 780–786 (2021).

54. K. Silliman, L. H. Spencer, S. J. White, S. B. Roberts, Epigenetic and Genetic Population Structure is Coupled in a Marine Invertebrate. Genome Biol. Evol. 15 (2023).

55. C. H. Waddington, Genetic assimilation of an acquired character. Evolution 7, 118–126 (1953).

56. M. J. West-Eberhard, Developmental Plasticity and Evolution (Oxford University Press, 2003).

57. M. J. Dubin, et al., DNA methylation in Arabidopsis has a genetic basis and shows evidence of local adaptation. Elife 4, e05255 (2015).

58. L. R. Feinberg, H. G. Dam, Effects of diet on dimensions, density and sinking rates of fecal pellets of the copepod Acartia tonsa. Mar. Ecol. Prog. Ser. 175, 87–96 (1998).

59. B. vonHoldt, E. Heppenheimer, V. Petrenko, P. Croonquist, L. Y. Rutledge, Ancestry-Specific Methylation Patterns in Admixed Offspring from an Experimental Coyote and Gray Wolf Cross. J. Hered. 108, 341–348 (2017).

60. F. Krueger, S. R. Andrews, Bismark: a flexible aligner and methylation caller for Bisulfite-Seq applications. Bioinformatics 27, 1571–1572 (2011).

61. B. Langmead, S. L. Salzberg, Fast gapped-read alignment with Bowtie 2. Nat. Methods 9, 357–359 (2012).

62. T. S. Jørgensen, et al., The Genome and mRNA transcriptome of the cosmopolitan calanoid copepod Acartia tonsa Dana improve the understanding of copepod genome size evolution. Genome Biol. Evol. 11, 1440–1450 (2019).

63. R Core Team, R: A Language and Environment for Statistical Computing. [Preprint] (2022). Available at: https://www.R-project.org/.

64. Y. Chen, B. Pal, J. E. Visvader, G. K. Smyth, Differential methylation analysis of reduced representation bisulfite sequencing experiments using edgeR. F1000Res. 6, 2055 (2017).

65. A. Alexa, J. Rahnenführer, T. Lengauer, Improved scoring of functional groups from gene expression data by decorrelating GO graph structure. Bioinformatics 22, 1600–1607 (2006).

66. A. Alexa, J. Rahnenfuhrer, Gene set enrichment analysis with topGO (2019).

67. M. Gautier, R. Vitalis, L. Flori, A. Estoup, ƒ□statistics estimation and admixture graph construction with Pool□Seq or allele count data using the R package poolfstat. Mol. Ecol. Resour. 22, 1394–1416 (2022).

68. A. R. Quinlan, I. M. Hall, BEDTools: a flexible suite of utilities for comparing genomic features. Bioinformatics 26, 841–842 (2010).

69. R. Kofler, R. V. Pandey, C. Schlötterer, PoPoolation2: identifying differentiation between populations using sequencing of pooled DNA samples (Pool-Seq). Bioinformatics 27, 3435–3436 (2011).

70. R. Patro, G. Duggal, M. I. Love, R. A. Irizarry, C. Kingsford, Salmon provides fast and bias-aware quantification of transcript expression. Nat. Methods 14, 417–419 (2017).

71. M. I. Love, W. Huber, S. Anders, Moderated estimation of fold change and dispersion for RNA-seq data with DESeq2. Genome Biol. 15, 550 (2014).

