## Supplemental Materials for "Complementary genetic and epigenetic changes facilitate rapid adaptation to multiple global change stressors"

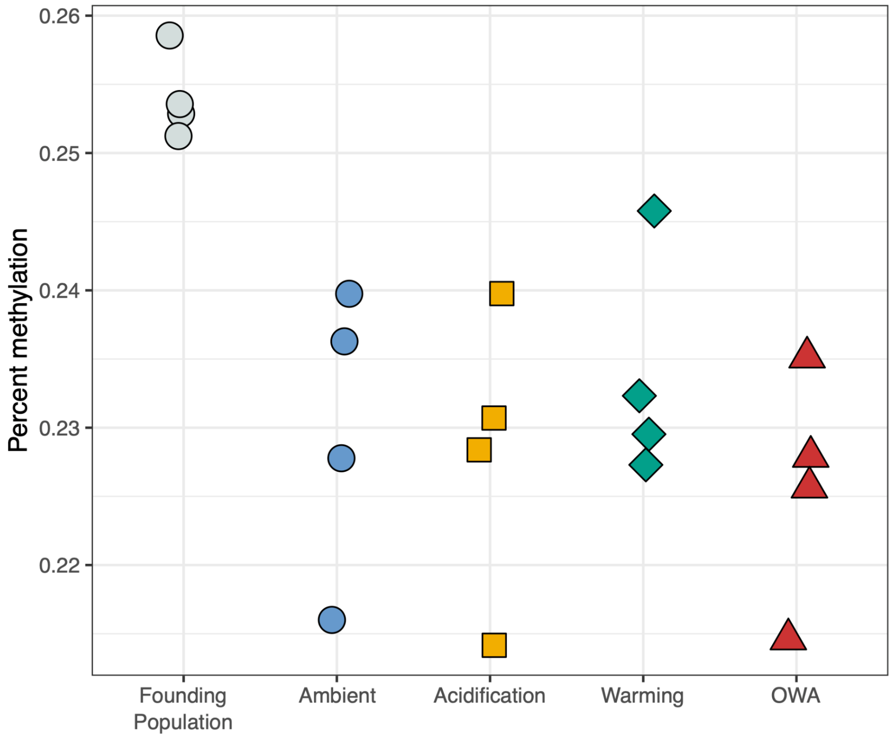


**Fig. S1.** Mean methylation percentage of all treatments. Each point is an individual replicate and color and shape indicate treatment.

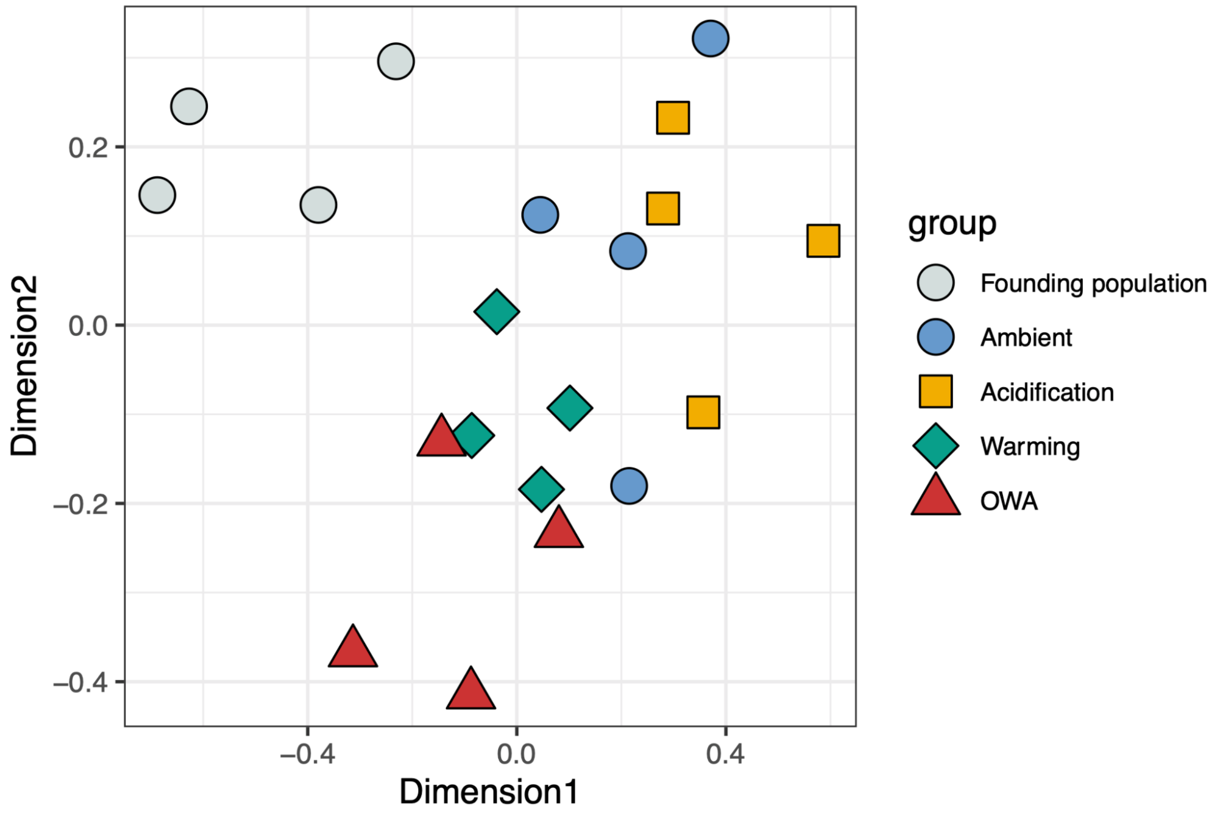


**Fig. S2.** Multidimensional scaling of methylation frequencies of 96,207 loci across replicate selection populations.

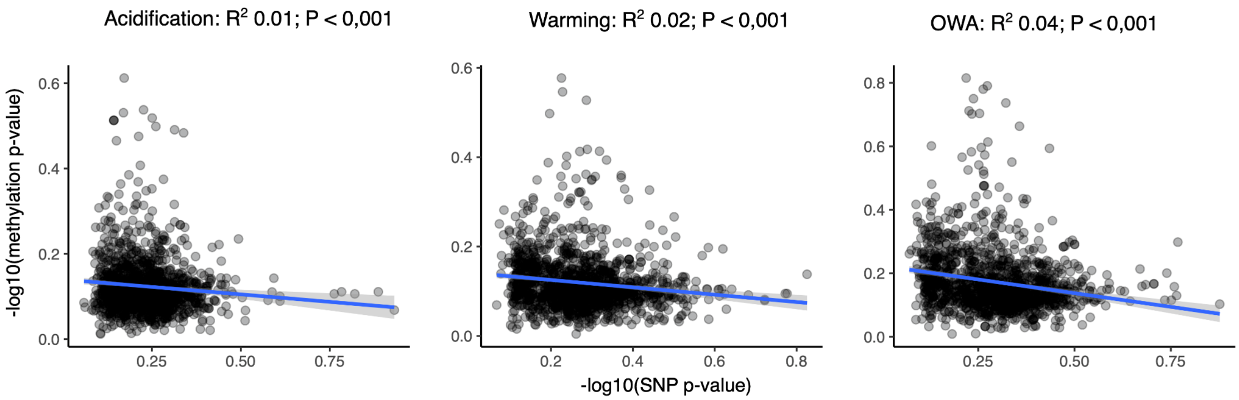


**Fig. S3:** 1349 genes with at least 4 SNPs and 4 methylation sites. This relationship holds if the number of snps/methylation sites required is changed.

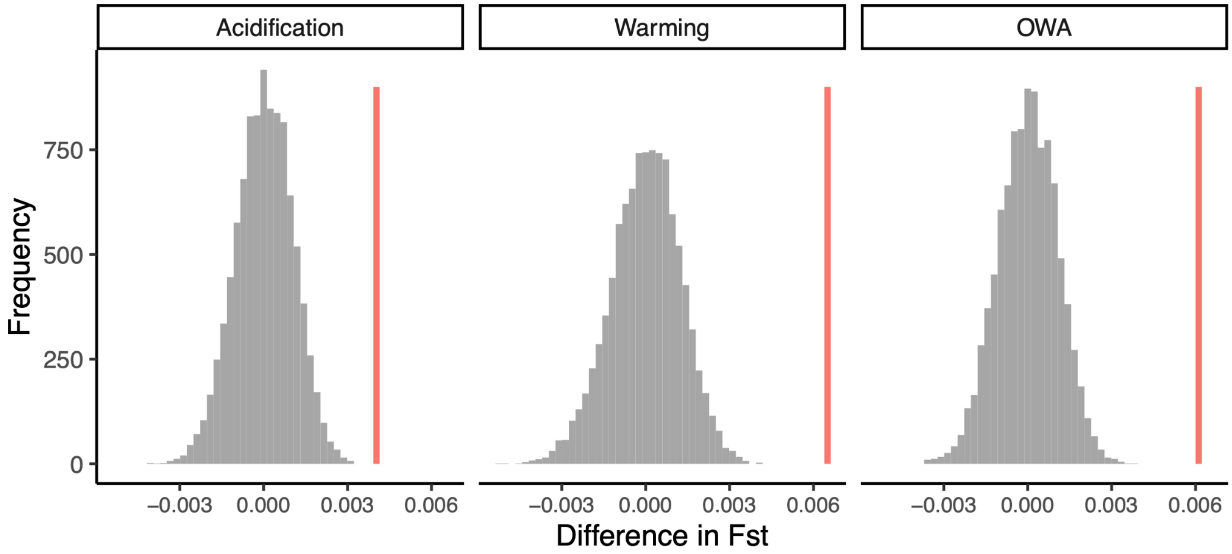


**Fig. S4:** Permutation test for the significance of the relationship between FST and methylation change. The genome was broken into 1.5 kb windows that contained at least 5 SNPs and 5 methylation sites resulting in 910 windows across the genome. The vertical pink line is the observed difference in mean $F_{ST}$ between windows containing significant versus no significant methylation site. To test if the observed difference in $F_{ST}$ was greater than expected by chance, a permutation test (n=10,000) was run, randomizing the windows categorized as significant for methylation change and the difference in $F_{ST}$ was calculated. Resulting p-values from these permutations: $p_{acidification} = 0.0001$; $p_{warming} = 0.0001$; $p_{OWA} = 0.0001$. This permutation corresponds to figure 3 in the main text.

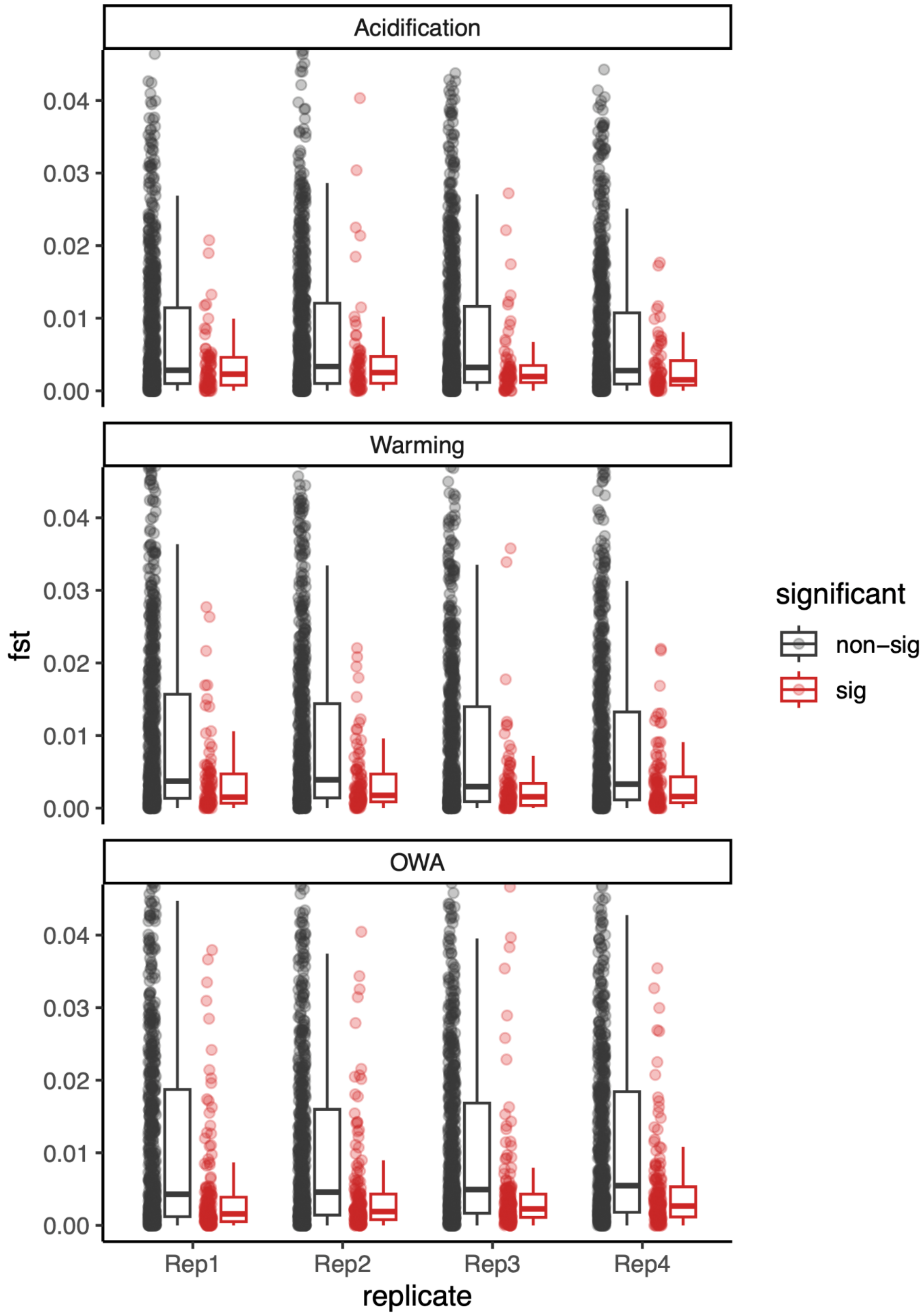


**Fig. S5:** Distribution of $F_{ST}$ values in 910 1.5 kb windows for each replicate and treatment. Windows contained at least 5 methylation and 5 SNP loci. Red windows ("sig") are those with at least one significant methylation change, black windows ("non-sig") contain no significant change.

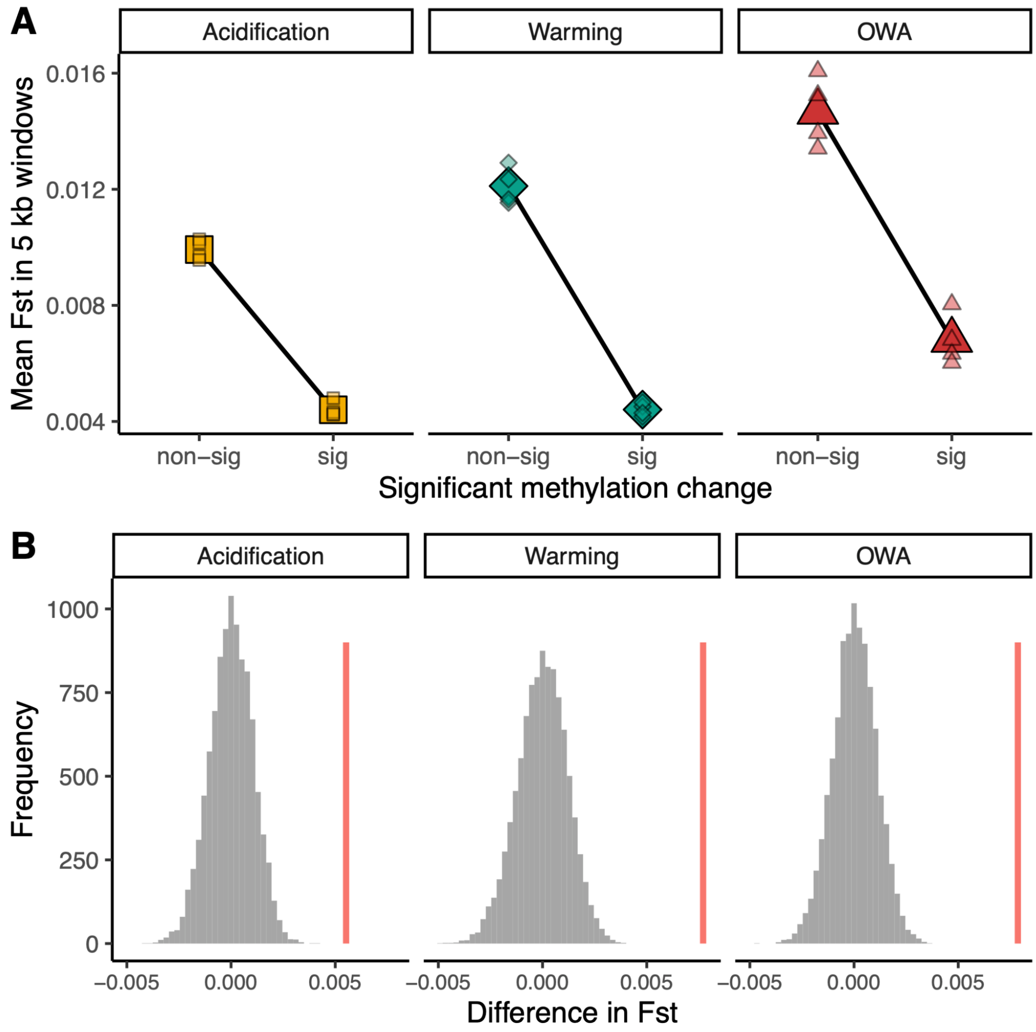


**Fig. S6:** Relationship between $F_{ST}$ and methylation change for acidification, warming, and OWA for 5kb window analysis; this analysis is the same as in Figure 2 but with a different window size. (A) The genome was broken into 5 kb windows that contained at least 5 SNPs and 5 methylation sites resulting in 1102 windows. Each small point represents a replicate and the large symbols are the mean of all replicates. If a window contained at least one significant change in methylation from the ambient treatment it was considered significant. (B) Permutation test (10,000) for the difference in $F_{ST}$ observed for each treatment. The vertical pink line is the observed difference in mean $F_{ST}$ between 5 kb windows containing significant versus no significant methylation site. Resulting p-values from these permutations: $p_{acidification} = 0.0001$; $p_{warming} = 0.0001$; $p_{OWA} = 0.0001$.

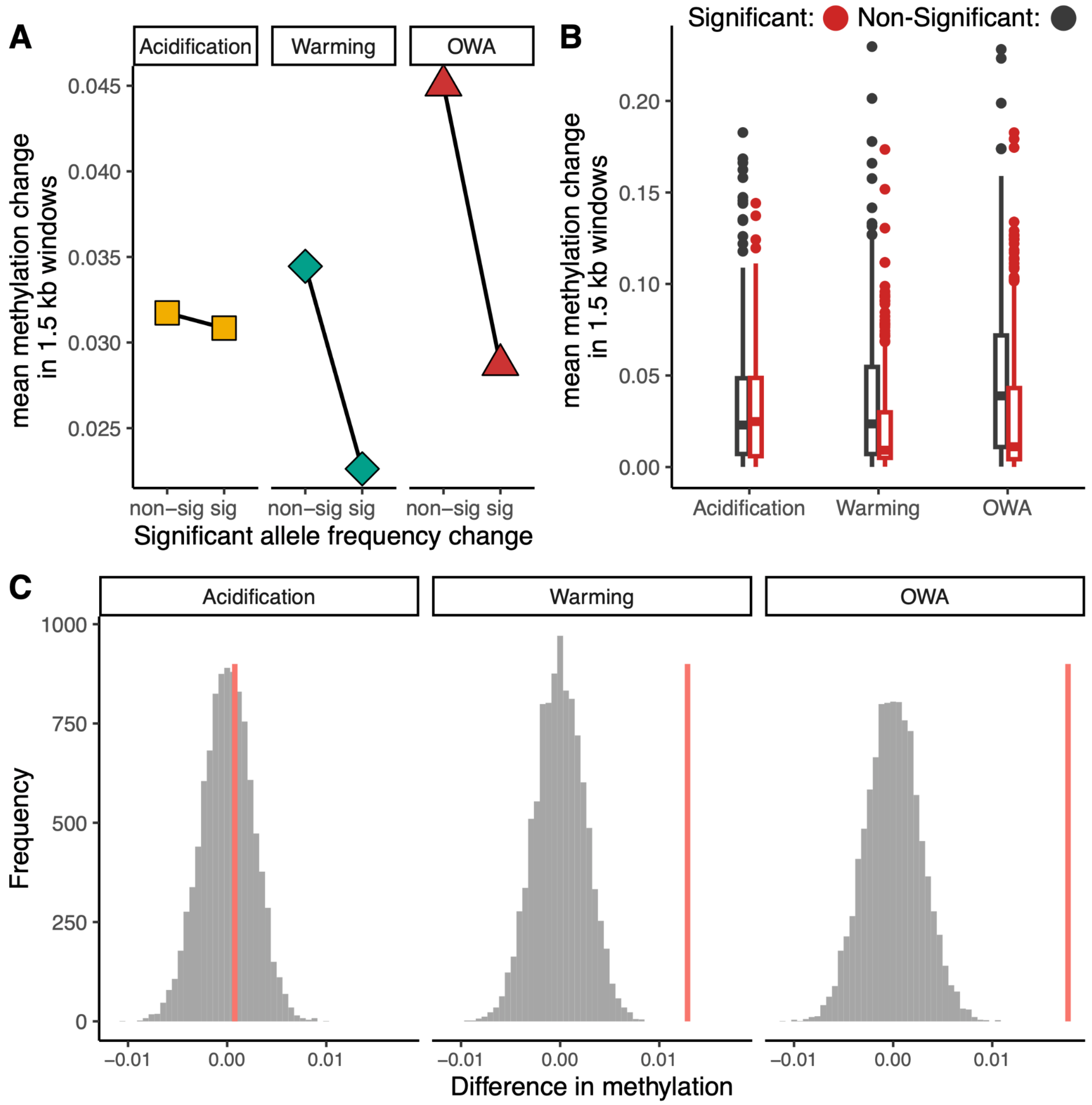


**Fig. S7:** Methylation changes for windows with significant allele frequency divergence. This is the opposite analysis from that presented in the main text. (A) The genome was broken into 1.5 kb windows that contained at least 5 SNPs and 5 methylation sites resulting in 910 windows across the genome. Each point represents the mean methylation percentage. If a window contained at least one significant change in allele frequency from the ambient treatment it was considered significant. (B) Distribution of mean methylation values shown in A. (C) Permutation test (10,000) for the difference in $F_{ST}$ observed for each treatment. The vertical pink line is the observed difference in mean $F_{ST}$ between 5 kb windows containing significant versus no significant methylation site. Resulting p-values from these permutations: $p_{acidification} = 0.3876$; $p_{warming} = 0.0001$; $p_{OWA} = 0.0001$.

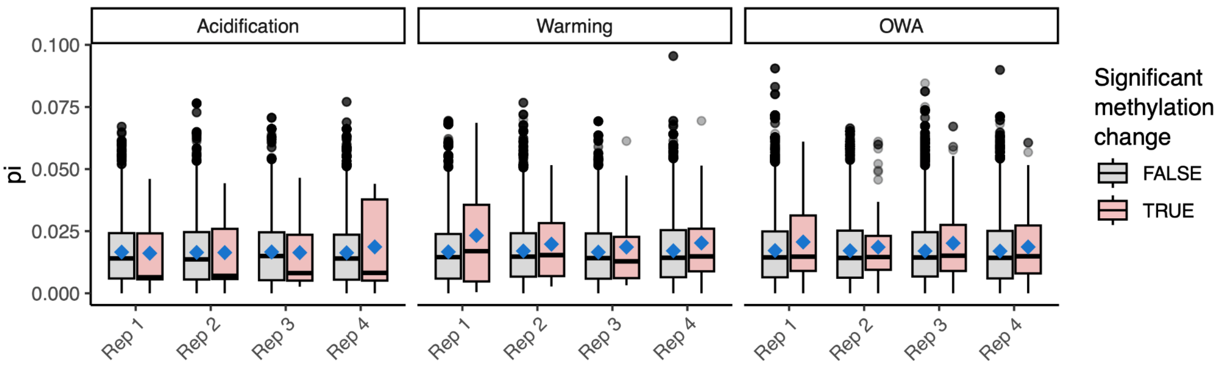


**Fig. S8:** Relationship between genetic variation and methylation changes. Tajima's π was calculated in 100 bp windows and all windows overlapping with a methylation loci were retained. Solid blue diamonds show the mean for each group and the horizontal lines show the median, as standard in a Tukey boxplot. For OWA, windows with significant methylation changes (TRUE) had significantly higher genetic diversity than those with no methylation change (FALSE; $p = 0.0001$; Kolmogorov-Smirnov Test). This pattern did not hold for acidification ($p = 0.2$) and warming ($p = 0.7$), though the number of windows overlapping with significant methylation changes was much lower than for OWA (acidification = 30; warming = 90; OWA = 456), which may lead to a poor ability to detect a signal.

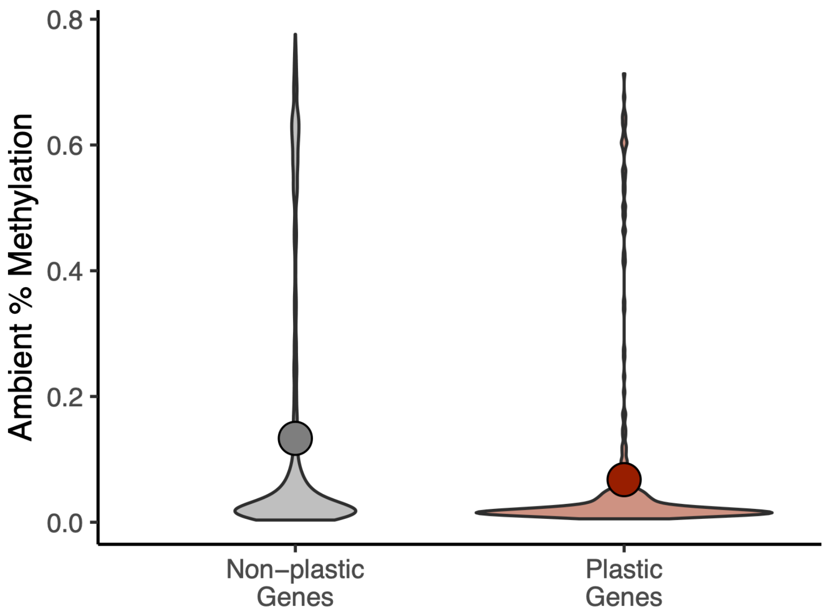


**Fig. S9:** Ambient plasticity genes vs. methylation. The percent methylation of plastic vs. non-plastic genes following transplant of the ambient line from ambient to OWA at generation F21. Solid points show the mean methylation percent for each group ( mean methylation ± se: Non-plastic: 0.133 ± 0.006; Plastic: 0.068 ± 0.007; mean ± se). Non-plastic genes had significantly higher methylation percent than plastic genes (p= 8.6e-8, Kolmogorov-Smirnov Test).

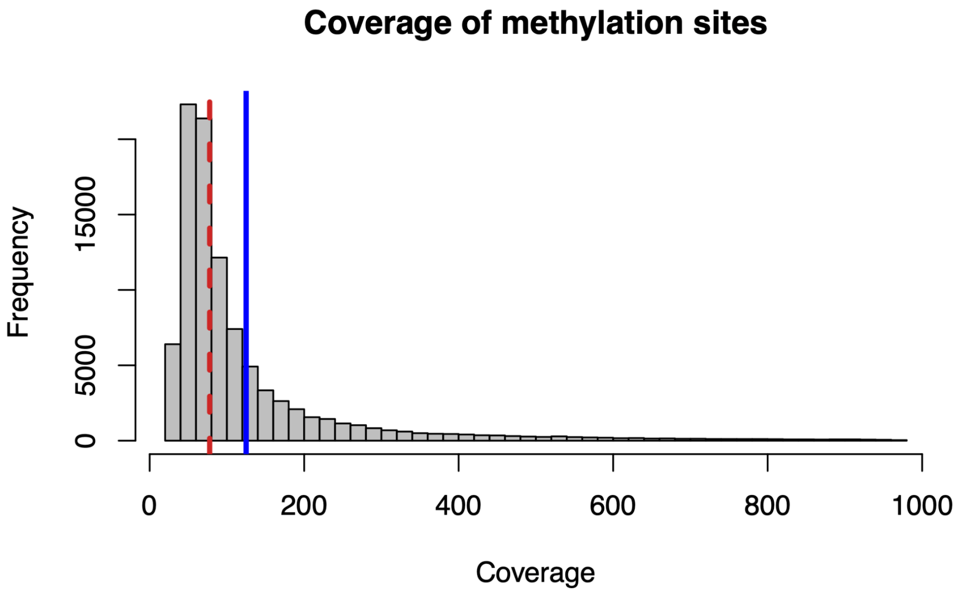


**Fig. S10:** Coverage histogram of methylated sites following filtering. The solid blue line is the mean coverage (124x) and the dashed red line is the median (77x).
